# DeepMap: A deep learning-based model with a four-line code for prediction-based breeding in crops

**DOI:** 10.1101/2023.07.26.550275

**Authors:** Ajay Kumar, Krishna T. Sundaram, Niranjani Gnanapragasam, Uma Maheshwar Singh, K. J. Pranesh, Challa Venkateshwarlu, Pronob J. Paul, Waseem Hussain, Sankalp Bhosale, Ajay Kohli, Berta Miro, Vikas Kumar Singh, Pallavi Sinha

**Affiliations:** International Rice Research Institute (IRRI), South-Asia Hub (SAH), Hyderabad, India; International Rice Research Institute (IRRI), South-Asia Regional Centre (SARC), Varanasi, India; International Rice Research Institute (IRRI), Metro Manila, Philippines; University of Missouri, Columbia, Missouri, USA

**Author notes:** Contributed equally (Ajay Kumar, Krishna T. Sundaram).

**Keywords:** DeepMap, Deep Learning, GPU, Quantitative phenotype prediction, Rice

## Abstract

Prediction of phenotype through genotyping data using the emerging machine or deep learning technology has been proven successful in genomic prediction. We present here a graphical processing unit (GPU) enabled DeepMap configurable deep learning-based python package for the genomic prediction of quantitative phenotype traits. We found that deep learning captures non-linear patterns more efficiently than conventional statistical methods. Furthermore, we suggest an additional module inclusion of epistasis interactions and training of the model on Graphical Processing Units (GPUs) in addition to Central Processing Unit (CPU) to enhance efficiency and increase the model’s performance. We developed and demonstrated the application of DeepMap using a 3K rice genome panel and 1K-Rice Custom Amplicon (1kRiCA) data for several phenotypic traits including days to 50% flowering (DTF), number of productive tillers (NPT), panicle length (PL), plant height (PH), and plot yield (PY). We have found that DeepMap outperformed the best existing state-of-the-art models by giving higher predictive correlation and low mean squared error for the datasets studied. This prediction performance was higher than other compared models in the range of 13-31%. Similarly for Dataset-2, significantly higher predictions were observed than the compared models (16-20% higher prediction ability). On Dataset-3, we have also shown the better and versatile performance of our model across crops (wheat, maize, and soybean) for yield and yield-related traits. This demonstrates the potentiality of the framework and ease of use for future research in crop improvement. The DeepMap is accessible at https://test.pypi.org/project/DeepMap-1.0/.

**Short Summary:** DeepMap is a deep learning-based breeder-friendly python package to perform genomic prediction. It utilizes epistatic interactions for data augmentation and outperforms the existing state-of-the-art machine/deep learning models such as Bayesian LASSO, GBLUP, DeepGS, and dualCNN. DeepMap developed for rice and tested across crops such as maize, wheat, soybean etc.

## INTRODUCTION

To be able to keep pace with the expected increase in food demand in the coming years, crop breeding must deliver the highest rates of genetic gains to maximize agricultural productivity. Deep Learning (DL) has emerged as a powerful tool in crop science, offering various applications such as predicting yield or quality traits from genotypes across different environments, plant disease recognition using Convolutional Neural Networks (CNNs), and image-based phenotyping using drones and edge computing devices (Albawi et al., 2017; Zou et al., 2019; Yaguchi et al., 2019). Harnessing the latent potential of DL’s non-linear and weighted architecture is a crucial step in leveraging DL for crop breeding applications. DL is a subset of Machine Learning (ML) methods that can identify complex patterns in large datasets and includes architectures such as Multi-Layer Perceptron (MLP), Deep Neural Network (DNN), Convolutional Neural Network (CNN), Recurrent Neural Network (RNN), Auto-encoders (AE), and Generative Adversarial Networks (GANs) (Goodfellow et al., 2014; McDowell et al., 2016).

Each DL architecture is designed for specific applications, with CNNs being particularly well-suited for image processing tasks such as object identification, document analysis, climate forecasting, medical image analysis, disease diagnosis, drug design, and protein structure prediction (Lundervold and Lundervold, 2019; Callaway, 2022; Renaud et al., 2021). RNNs, on the other hand, excel in handling sequential and temporal data, such as text or videos. Deep Neural Networks offer the ability to capture additional input features, while GANs and AE are used to generate new data from existing examples, increasing the sample size and improving model accuracy and performance. DL algorithms, as non-parametric methods, are more efficient in identifying non-linear patterns compared to traditional genome-based machine learning methods (Pratley, 2003; Pérez-Rodríguez et al., 2012; Montesinos-López et al., 2021; van Dijk et al., 2021; Li et al., 2021).

DL provides the flexibility to map complex associations between data and output, relying on high-quality and sufficiently large training data. Over the past decades, various pattern recognition models have been employed for genotype-to-phenotype prediction, including Bayesian Artificial Neural Networks (BNNs), Regularized Neural Networks, Deep Belief Network (DBN), Reproducing Kernel Hilbert Space (RKHS), Bayesian LASSO (B-LASSO), Best Linear Unbiased Prediction (BLUP), and deep convolutional neural network (dCNN) (Gianola et al., 2011; Rachmatia et al., 2017; Ma et al., 2018). Ensemble methods, such as the Stacking Ensemble Learning Framework (SELF) and combinations of statistical techniques and machine learning models, have been shown to improve model performance (Guzzetta et al., 2010; Endelman 2011, Liang et al., 2021; Munneb and Henschel, 2021).

Deep neural networks have the advantage of being able to learn from millions of data points without reaching a performance plateau. This computational tractability is achieved by leveraging accelerators such as Graphical Processing Units (GPUs), Tensor Processing Units (TPUs), and Information Processing Units (IPUs), as well as parallel file system technologies (Renaud et al., 2021). While genomic prediction (GP) methods have been developed and made available through scripts and CRAN packages, there is a need to optimize these algorithms with evolving deep learning techniques for the ease of use by non-coding communities. Thus, the development of a user-friendly Python package named DeepMap, consisting of just four lines of code, would be highly beneficial for both core researchers and interdisciplinary communities.

In this manuscript, we describe the structure of the DeepMap framework and demonstrate its applicability and potential for genomic prediction using three rice datasets. Our results show improved accuracy in terms of Pearson correlation on Dataset-1 (IRRI-SAH) by 9-30%, on Dataset-2 (IRRI-SARC) by 11-26%, and on Dataset-3 (1kRiCA) by 23-33% compared to existing state-of-the-art ML and DL models. Furthermore, DeepMap has the versatility to be applied across different crops and for both qualitative and quantitative traits.

## RESULTS

### Description of DeepMap

DeepMap is a deep learning-based prediction model built with a python3^21^ that allows end-to-end model training and prediction of quantitative phenotypic trait value from the genotyping data. The four-line code takes genotypic and phenotypic data in the required format. Then, it calls the main function to train the model and give the output in comma-separated files for ‘k’ cross-validations of predicted values, model training/validation loss, and scatter plot of actual vs. predicted plots. Conceptually, the SNPs information along with the epistatic interaction is passed to the fully connected seven layer deep neural network architecture that leads to the single neural unit output of predicted phenotypic trait value. The parameters can be changed and passed through a function call which gives flexibility to the function and gives a scope of optimization. The overall workflow from data generation to model prediction can be found in **Figure 1**.

**Figure 1.**
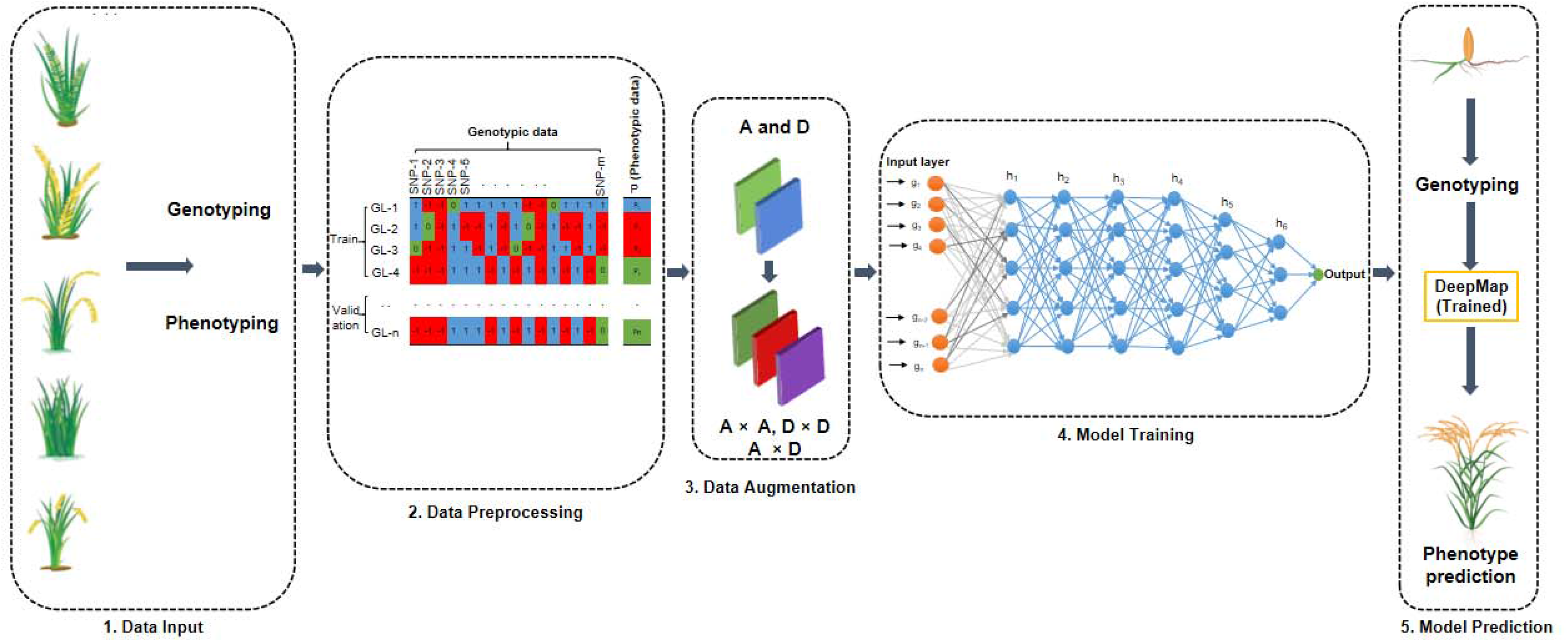
The DeepMap framework. The DeepMap framework is divided into continuous step processes from data generation to model prediction. 1) data input. Starting from genotyping and phenotyping of anticipated crop and observed phenotyping trait of interest for mapping. 2) data pre-processing. Organized genotyping and phenotyping data and convert alleles into ‘0’ representing homozygous allele-2, ‘1’ is representing heterozygous allele and ‘-1’ is representing homozygous allele-1/missing alleles for n genotyping lines (GL). Further, data converts into additive and dominance interactions. 3) data augmentation. In this step, epistasis interactions are augmented using additive and dominance matrix information provided along with the SNPs data as an input to the model. The additive interactions (AI) and dominance interactions (DI) are used to generate additive-additive interactions (AI×AI), dominance-dominance interactions (DI×DI), and additive-dominance interactions (AI×DI). These five epistatic interactions are used to train the model. 4) model training. The training dataset is given to the model for learning genomic patterns. For each genotypic line, G^i^ = {g_1_, g_2_, g_3,…,_ g_m_} where ‘m’ is the number of epistatic interactions amid genotypic lines for ‘n’ SNPs. As per the phenotypic trait of interest, the predicted output might be quantitative or qualitative. The proposed model can be used for the qualitative trait of interest by changing the output function to sigmoid or SoftMax activation function in the output layer. 5) model prediction. The validation dataset is given to the model to check the performance of the model on an unseen dataset. The trained model can be used to predict phenotypic trait value prediction by giving SNPs and epistatic interactions to the trained model. The performance of the model could be optimized by changing the hyperparameters of the model. **Abbreviations:** SNPs: single nucleotide polymorphism; GL: genotypic line; AI: additive interactions; DI: dominance interactions; AI×AI: additive-additive interactions; DI×DI: dominance-dominance interactions; AI×DI: additive-dominance interactions; h: hidden layer.

The framework consists of five major steps, which are as follows:

#### Data input

The phenotypic values are generated for the trait of interest after following the experimental design. The corresponding genomic information can be generated through either sequencing (whole genome, exome sequencing) or genotyping (GBS, SNP chip). Since most significant crops are sequenced and available in open source, the genomic information can be downloaded from their respective databases. The genotypic data and its corresponding phenotyped trait values are passed to the model for prediction.

#### Data preprocessing

As per raw genotypic and phenotypic data comprehends an ample amount of manual (human) and field errors. Therefore, data pre-processing is one of the major steps to clean up the data by removing missing values and unexpected observations. The genotypic data should be cleaned for Minor Allele Frequency (MAF), missingness percentage, and can be reduced further based on the LD (Linkage Disequilibrium) or co-linearity between the markers. Further, conversions are performed on genotypic data to convert the raw data into model-required format (converting plink file (PLINK 2.00 alpha accessed through link: https://www.cog-genomics.org/plink/2.0/) format into .bed (PLINK binary biallelic genotype table), .bim (PLINK extended MAP file), and .fam (PLINK sample information file) files). The proposed model requires four input files that include, three files of marker genotypic data (Single Nucleotide Polymorphisms (SNPs) data in numerical format **(Table S1)**, additive **(Table S2)**, and dominance (**Table S3**) interactions) and a single file of observed phenotypic trait data **(Table S4)**. **Figure 1** shows a dataset of ‘n’ number of Genotypic Lines (GL) with ‘m’ number of SNPs. SNPs encoded as ‘0’ represent homozygous allele 2, ‘1’ represents heterozygous allele, and ‘-1’ represents homozygous allele 1.

The ‘-1’ was also represented for missing alleles, as the frequency was very low in the dataset. The observed phenotypic trait values were passed as a single column [P = {P_1_, P_2_, P_3_,…., P_n_}, for each GL] through the model. The processed dataset was then divided into two groups called training set (80%) and validation set (20%) using the K-fold algorithm of sklearn (Garreta, 2013) python package (percentage of train and validation set can be altered while calling the function for each dataset). Subsequently, the training set genotypic and phenotypic data undergo a downstream pipeline to train the model, and the validation set is unseen to the model which we use to predict the phenotypes and correlates with actual phenotypic trait values of the validation set.

#### Data augmentation

In this step, epistatic interactions are augmented using additive and dominance matrix information provided along with the SNPs data as an input to the model. In **Figure 1**, the additive information (A) and dominance information (D) are used to generate additive-additive interactions (A×A), dominance-dominance interactions (D×D), and additive-dominance interactions (A×D). These five epistatic interactions are used to train the model.

#### Model training

The training dataset contains both the phenotypic and genotypic data given to the model for learning the genomic patterns corresponding to the phenotypic trait value. For i^th^ genotypic line, GL^i^ = {*g*1 *i*, *g*2 *i*, *g*3 *i*, …, *gn i*} where *gn i* is the epistatic interaction of i^th^ and n^th^ genotypic line, and ‘n’ is the number of genotypic lines. The model was trained using a deep neural network algorithm and the hyperparameter (based on grid search; further information is available in the methods section) is optimized to increase the model’s performance and reliability. As per the phenotypic trait of interest, the predicted output might be quantitative or qualitative. In DeepMap, we have used ten-cross validations for quantitative complex phenotypic traits for the prediction. The proposed model can be used for the qualitative trait of interest by changing the output function to a sigmoid or softmax activation function in the output layer.

#### Model prediction

The validation dataset contains only genotypic information and is given to the model to check the model’s performance by comparing the predicted phenotypic value with the actual value. The performance of the model could be optimized by changing the hyperparameters of the model.

### Evaluation of model performance

We independently trained the model on three different rice datasets (Dataset-1, Dataset-2, and Dataset-3). To check the versatility of our model to use for other crops, we trained and validated DeepMap with datasets of wheat, maize, and soybean. For the performance evaluation, we compared the predictive ability of DeepMap with Bayesian LASSO, rrBLUP, DeepGS, and dualCNN in all the selected data sets.

### DeepMap application in Dataset-1

A set of 2,229 rice varieties from 3K rice panel (3K RGP, 2014) phenotyped at International Rice Research Institute – South Asia Hub (IRRI-SAH) (Hyderabad, India), have been used as the first dataset to train and evaluate DeepMap. We selected five yield and yield related traits (DTF, NPT, PL, PH, and PY) because they showed significant variation among themselves and had higher heritability, making them highly suitable for GP (**Table 1**).

**Table 1.** Descriptive statistics of 3K rice accessions phenotyped in two locations.

| Datasets | Trait | No of lines | H<sup>2</sup> | Mean<sup>*</sup> | SD<sup>*</sup> |
| --- | --- | --- | --- | --- | --- |
| Dataset-1<br>IRRI-SAH<br>(27.2046° N, 77.4977° E) | DTF | 532 | 0.89 | 96.87 | 11.46 |
|  | NPT | 1,316 | 0.82 | 10.44 | 2.32 |
|  | PL | 1,223 | 0.77 | 19.24 | 1.73 |
|  | PH | 963 | 0.91 | 111.17 | 19.22 |
|  | PY | 1,163 | 0.72 | 209.15 | 87.66 |
| Dataset-2<br>ISARC<br>(25.3024° N, 82.9491° E) | DTF | 532 | 0.88 | 87.62 | 15.70 |
|  | NPT | 1,316 | 0.76 | 7.89 | 1.34 |
|  | PL | 1,223 | 0.83 | 24.09 | 2.54 |
|  | PH | 963 | 0.85 | 141.94 | 21.94 |
|  | PY | 1,163 | 0.71 | 254.40 | 91.85 |

**Abbreviations:** PY: plot yield (gram); PH: plant height (cm); PL: panicle length (cm); NPT: number of productive tillers; DTF: days to 50% flowering (number of days); N<sup>a</sup>: number of phenotyped rice varieties; H<sup>2</sup>: broad-sense heritability; SD: standard deviation. IRRI: International Rice Research Institute, ISARC: IRRI South Asia Regional Centre.
\*Mean and SD are calculated on BLUPs.

The experiment was performed on DeepMap for ten-cross validations on these five phenotypic traits of IRRI-SAH location (Supplementary **Table S5** to **Table S14** for prediction performance and cross-validation results). Average Pearson Correlation Coefficient between the predicted and observed phenotypic values for the validation dataset was 0.74, 0.65, 0.67, 0.76, and 0.70 for DTF, NPT, PL, PH, and PY (**Figure 2a)**, respectively. The DeepMap outperformed (**Table 2**) standard best-performing methods by 14%, 19%, 25%, 13%, and 31% for the DTF, NPT, PL, PH, and PY traits, respectively. In **Table 3**, states the metric of model evaluation for model performance where, Mean Squared Error (MSE) is 0.52, 0.71, 0.67, 0.47, and 1.98 on DTF, NPT, PL, PH, and PY respectively (**Figure 2d)**, obtained on 10,000 epochs. As there is no upper limit of MSE, but the lower MSE represents the lower error in predicted phenotypic trait values. Thus, minimized error (less than one) in DTF, NPT, PL, PH shows significant predictions, and MSE > 1 in PY shows that phenotypic trait is complex to predict, where MSE can be further reduced by hyperparameter optimization. The stretched bound of Pearson correlation and MSE of NPT shows the diversity in mapping and complexity of prediction. The Coefficient of determination metric is an additional statistical measure for the regression model and the corresponding scoring for DTF, NPT, PL, PH, and PY are 0.49, 0.03, 0.24, 0.55, and 0.43, respectively. In **Figure S1**, the training and validation dataset shows the convergence of training and validation (testing) loss that substantiates the model’s prediction ability. The training and validation loss converges at the 1433^rd^ epoch for DTF, 148^th^ epoch for NPT, 587^th^ epoch for PH, 377^th^ epoch for PL, and 3987^th^ epoch for PY. Comparatively, plot yield (PY) takes more iteration and time to converge the losses. The predicted vs actual value graph (**Figure S2**) reveals the prediction performance of the model.

**Figure 2.**
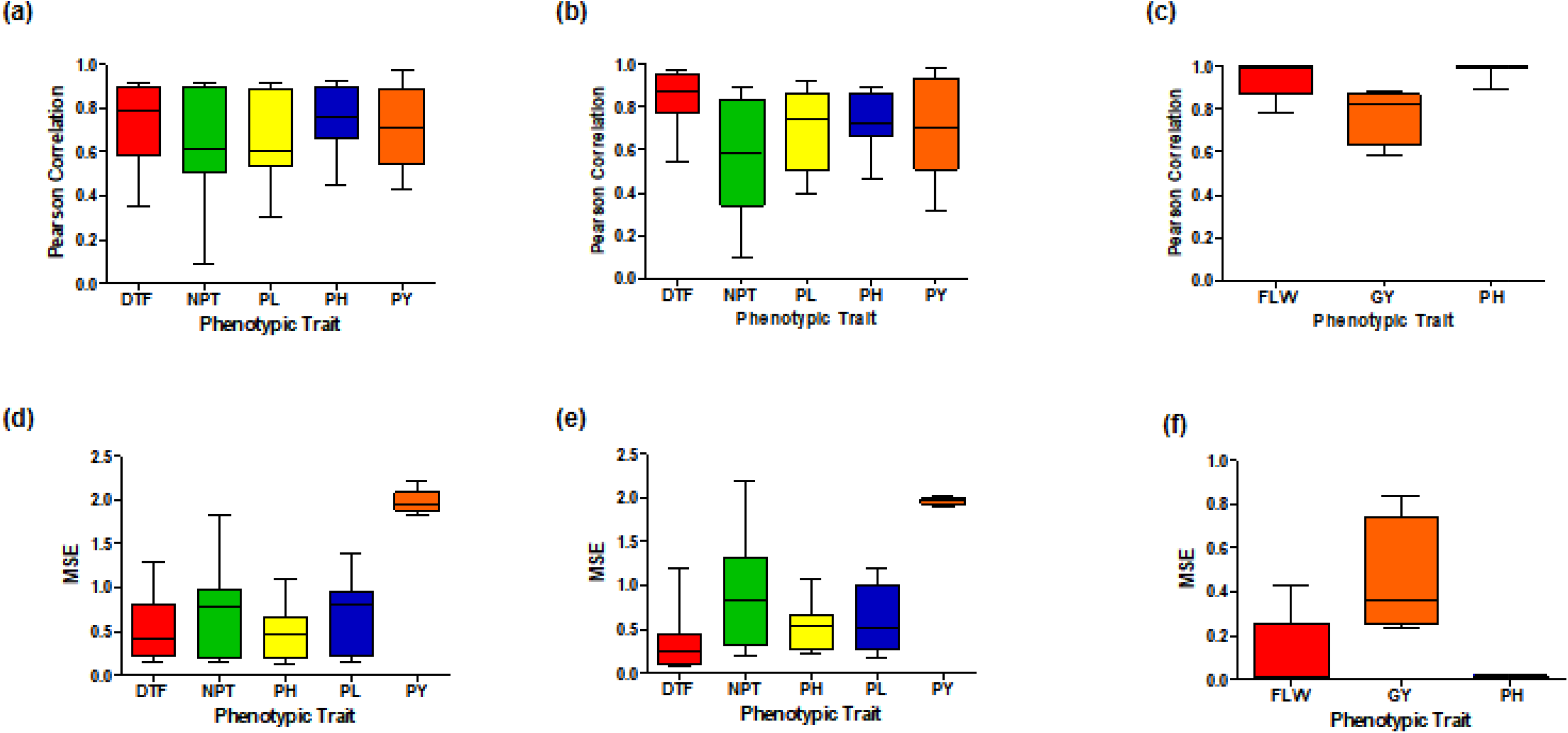
Pearson correlation and Mean Squared Error (MSE) on case datasets. a) Pearson correlation of ten-cross validation for five phenotypic traits DTF, NPT, PH, PL, PY on Dataset-1 IRRI-SAH. b) Pearson correlation of ten-cross validation for five phenotypic traits DTF, NPT, PH, PL, PY on Dataset-2 ISARC. c) Pearson correlation of ten-cross validation for three phenotypic traits FLW, GY, PH on Dataset-3 1kRiCA. d) mean squared error (MSE) of ten-cross validation for five phenotypic traits DTF, NPT, PH, PL, PY on Dataset-1 IRRI-SAH. e) MSE of ten-cross validation for five phenotypic traits DTF, NPT, PH, PL, PY on Dataset-2 ISARC. f) MSE of ten-cross validation for three phenotypic traits FLW, GY, PH on Dataset-3 1kRiCA. **Abbreviations:** DTF-days to 50% flowering (number of days); FLW: flowering time (number of days for flowering); NPT-number of productive tillers (numbers); PH-plant height (cm); PL-panicle length (cm); PY-plot yield (kg/hectare); GY-grain yield (gram/plot).

**Figure 3.**
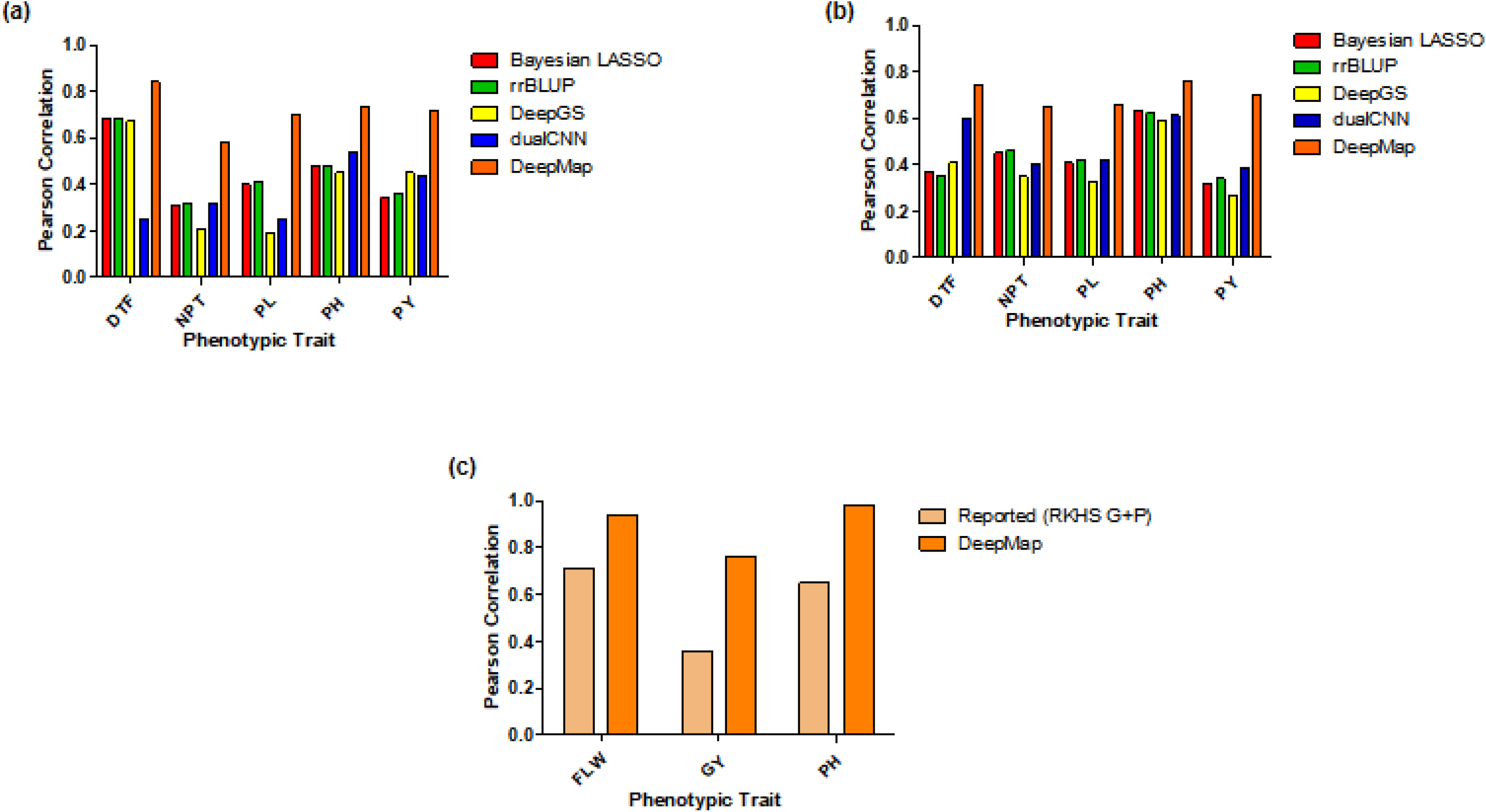
Model comparison. a) The five phenotypic traits of Dataset-1 IRRI-SAH Pearson correlation is compared among DeepMap and standard ML-DL methods, including Bayesian-LASSO, rrBLUP, DeepGS, and dualCNN. b) The five phenotypic traits of Dataset-2 ISARC Pearson correlation is compared among DeepMap and standard ML-DL methods, including Bayesian-LASSO, rrBLUP, DeepGS, and dualCNN. c) The three phenotypic traits of Dataset-3 1kRiCA pearson correlation are compared with the performance of DeepMap and the best-reported model (RKHS G+P). **Abbreviations:** ML: machine learning; DL: deep learning; dualCNN: dual convolutional neural network; DeepGS: deep genomic selection; rrBLUP: ridge-regression bayesian linear unbiased prediction; LASSO: least absolute shrinkage and selection operator. RKHS: reproducing kernel hilbert space. 1kRiCA: 1K-Rice custom Amplicon; IRRI-SAH: International Rice Research Institute (IRRI) South Asia Hub; ISARC- IRRI South Asia Regional Centre.

**Table 2.**
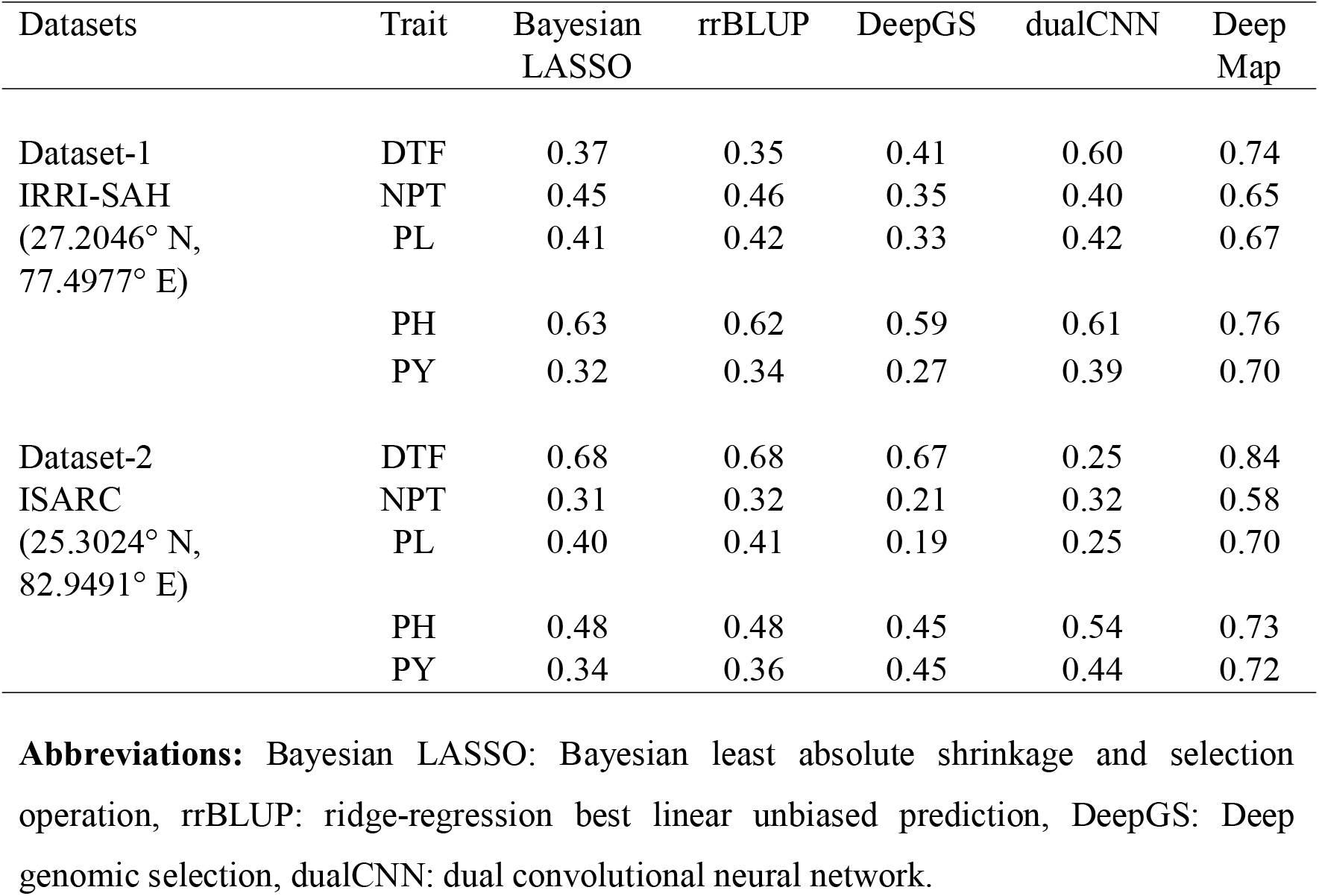
Pearson correlation coefficient of five models performed for two datasets for five phenotypic traits. Fitted models were Bayesian LASSO, rrBLUP, DeepGS, dualCNN, and DeepMap. The results are performed on ten cross-validations for all models and the correlation coefficient is observed on unseen genotyped rice lines to the trained model.

**Table 3.**
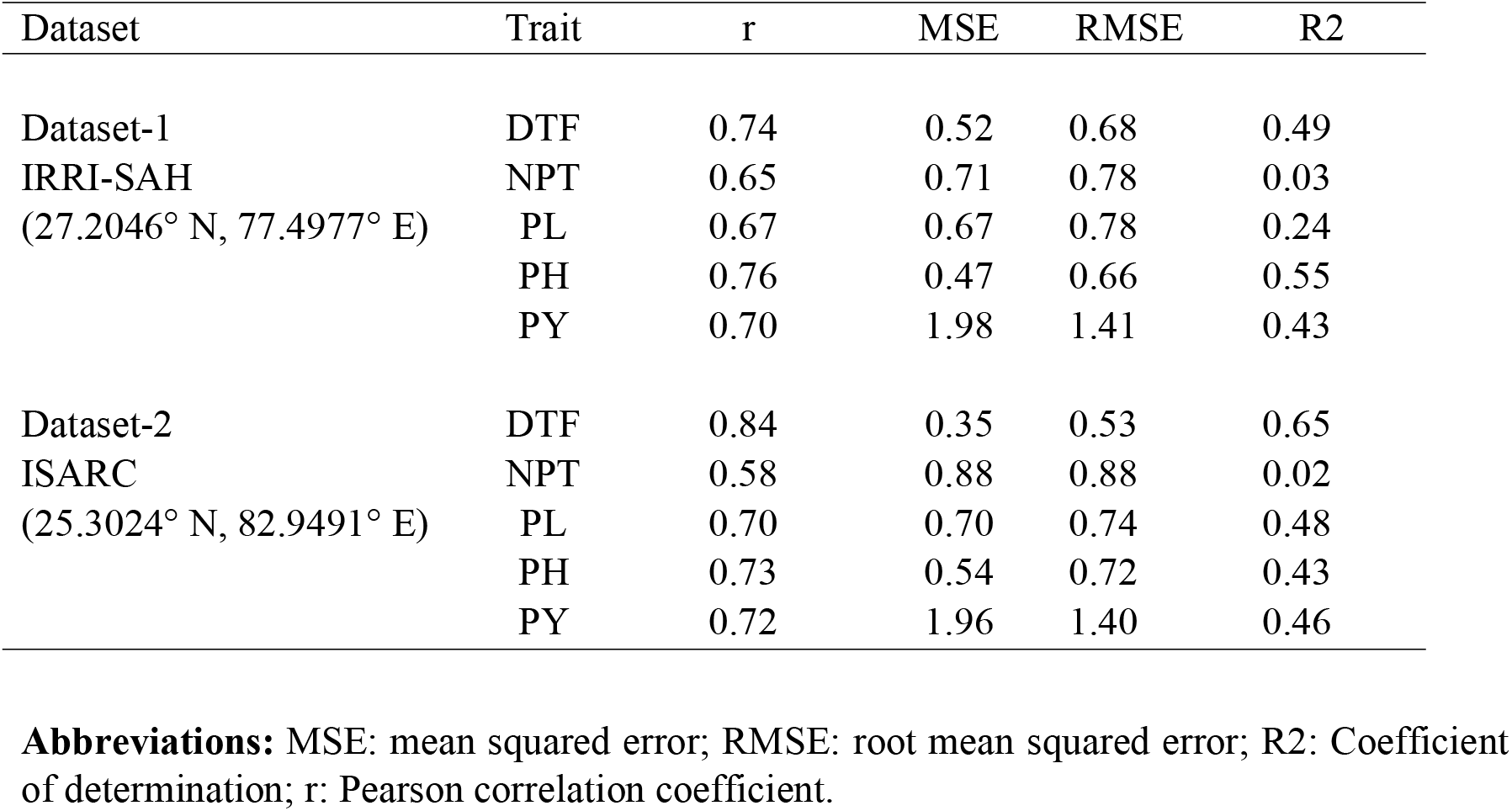
Model evaluation parameters Pearson Correlation Coefficient, mean squared error, root mean squared error, and Coefficient of determination is calculated on the validation dataset (unseen to the model) for discerning the performance of the model.

### DeepMap application in Dataset-2

A set of 2,145 rice varieties from 3K rice panel (3K RGP, 2014) phenotyped at International Rice Research Institute - South Asia Regional Centre (ISARC) (Varanasi, India), have been used as the dataset 2 for the model training and evaluation. The same yield and yield related traits as in Dataset-1 were used here as well (**Supplementary Table S15** to **Table S24** for prediction performance and cross-validation results). An average Pearson Correlation Coefficient (r) were found to be 0.84, 0.58, 0.70, 0.73, and 0.72 for DTF, NPT, PL, PH, and PY, respectively (as shown in **Figure 2b)** in the unseen/validation dataset. The DeepMap outperformed (Shown in **Table 2**) standard best-performing methods by 15%, 25%, 29%, 19%, and 27% for the DTF, NPT, PL, PH, and PY traits. **Table 3** states the metric of model evaluation for model performance where, Mean Squared Error (MSE) is 0.35, 0.88, 0.70, 0.54, and 1.96 on DTF, NPT, PL, PH, and PY, respectively (as shown in **Figure 2e**), obtained on 10,000 epochs. Thus, minimized error (less than one) in DTF, NPT, PL, and PH shows significant predictions, and MSE > 1 in PY shows that phenotypic trait is complex to predict, where MSE can be further reduced by hyperparameter optimization. The stretched bound of Pearson correlation and MSE of NPT phenotypic trait shows the diversity in mapping and complexity of prediction. The Coefficient of determination for the regression model is 0.65, 0.02, 0.48, 0.43, and 0.46 for DTF, NPT, PL, PH, and PY, respectively. As seen in Dataset-1, (the prediction ability of our model is high for this dataset too, which was validated through the convergence of training and validation (testing) loss (**Figure S1**). The training and validation loss converges at the 864^th^ epoch for DTF, 206^th^ epoch for NPT, 945^th^ epoch for PH, 3488^th^ epoch for PL, and 6245^th^ epoch for PY. Since we have chosen the common germplasm for all the traits in Datasets-1 and -2, the results showed a similar trend for the traits. The complex phenotypes such as PY are difficult to predict, comparatively. The predicted vs actual value graph (**Figure S2**) reveals the prediction performance of the model for Dataset-2.

### DeepMap application in Dataset-3

This dataset consists of 353 accessions genotyped using the 1K-Rica (Arbelaez et al., 2019) Custom Amplicon (1k-RiCA), a robust custom sequencing-based amplicon of 967 SNPs. We considered three phenotypic traits, flowering time, grain yield, and plant height to calculate the predictive ability of DeepMap. The reported predictive abilities of the best performing model (RKHS G+A) based on genomic selection were 0.71, 0.36, and 0.65 for Flowering time, grain yield, and plant height, respectively (as shown in **Figure 2c)**. The DeepMap reported 0.94, 0.76, and 0.98 for Flowering time, grain yield, and plant height, respectively, and outperformed the state-of-the-art genomic prediction models in the range of 23-40% (Supplementary **Table S25** to **Table S30** for prediction performance and cross-validation results). **Table S25** states the metric of model evaluation for performance where the Mean Squared Error (MSE) is 0.11, 0.49, and 0.01 on the corresponding phenotypic traits as shown in **Figure 2f**.

### Validation of DeepMap across crops

We have also performed the genomic prediction across crops including wheat, maize, and soybean. The 1275 genotypes of wheat were considered with 5741 SNPs for spike grain number (SGN) and time young microspore (TYN). The Pearson correlation for SGN and TYM obtained was 0.48 and 0.85 (9-12% increase with DeepMap). The 309 genotypes of maize were used with 309 SNPs for DTF and Grain Yield (GY). The predictive ability achieved was 0.54 and 0.73 (14-19% increase). The soybean’s 5558 genotypic lines with 5140 SNPs for phenotypic traits including yield, protein and height were considered. The Pearson correlation achieved was 0.56, 0.80 and 0.86 (11-24% increase with DeepMap).

## DISCUSSION

We have presented our DeepMap framework, demonstrating its uses and performance on three case study datasets. Its main advantages are as follows:

1. From the user’s perspective, DeepMap is hosted on Python Package Interface (PyPI accessed using link https://pypi.org/) (DeepMap PyPI link is in code availability section) and the four-line-code execution of the pipeline in a single program run makes it easy to use, unlike previously available lengthy R scripts. It works as a black box for the non-coding communities and, gives a scope of hyper parameterization to increase the performance by changing the function call parameters. The generalized architecture of DeepMap for genotype to phenotype prediction makes it directly applicable to a range of genomic selection challenges.
2. From a developer’s perspective, DeepMap is an open-source deep learning-based python package with version control, continuous integration, and easy addition of new features hosted on GitHub (Link is available in code availability section). This flexibility increases the maintainability and future developments in DeepMap by the community, for example, the addition of crop data like maize, wheat, and Soybean.
3. Computational efficiency: DeepMap is developed to leverage Graphical Processing Units (GPUs) for faster training and better prediction accuracy. However, it can also work on Central Processing Units (CPUs) but it would take more training time. The future of DeepMap’s computational power is to deploy on Amazon Web Services (AWS) and utilize cutting-edge hardware accelerators like GPUs, TPUs, and IPUs.
4. Finally, the competing performance and ability to outperform the state-of-the-art methods (**Table 2**) demonstrate that the utilization of a non-linear deep learning framework and inclusion of epistasis interactions improves the model’s prediction ability.

In summary, DeepMap is the first python package based on deep learning for genotype to phenotype prediction. It features a four-line code for genomic selection for quantitative phenotypes. Compared with other well-known machine and deep learning methods (Bayesian methods, GBLUP (Tuberosa and Crossa, 2019), rrBLUP, and DeepGS), DeepMap shows higher prediction accuracy for the quantitative phenotypic prediction (**Table 2**). The DeepMap can also be employed for qualitative (categorical) phenotype prediction by changing the output activation function in the last layer. The accuracy of the model can be increased by developing region-based (location-based) or variety-based (crop/species-based) prediction models that would open doors to widen genomic prediction challenges.

In the future, we plan to extend our model’s scope by incorporating multiple input factors such as environmental (Montesinos-López et al., 2018) information, soil information, and image-based phenotyping data. We will expand the package to support automated deep learning (AutoDL) to automatically adjust hyperparameters for ease of use for non-coding communities and extend the use of prediction-based breeding across crops/species. These features will be available in the upcoming version of DeepMap. We have used epistatic interactions that shows our model is working better than the standard machine/deep learning models. We have used our model on various crops like maize, wheat, and soybean that shows that it is applicable to other crops also. We have used three datasets of rice of different locations that show it can be used for different environments/ locations.

## MATERIALS AND METHODS

### Generation of plant datasets

Two of the datasets (Dataset-1 & Dataset-2) used in our study are from 3K rice panels. The Dataset-1 consists of 2,229 rice accessions and had been evaluated at International Rice Research Institute – South Asia Hub (IRRI-SAH), Hyderabad, India (17.4993° N, 78.2759° E). Dataset-2 is a set of 2,145 rice accessions and had been evaluated at International Rice Research Institute - South Asia Regional Centre (ISARC), Varanasi, India (25.3024° N, 82.9491° E). Both subsets were grown under irrigated-Transplanted Rice (TPR) condition during wet season-2019 (commenced on 18th July 2019) with an objective to evaluate various yield and yield related traits. The plants were organized in an Augmented Randomized Complete Block Design (RCBD) with four checks repeated in each block.

The selected five traits (DTF, NPT, PL, PH, and PY) were analyzed individually using residual maximum likelihood (REML) in GenStat 17 (https://www.vsni.co.uk/) in a mixed model approach considering genotypes as random effect and block as a fixed effect. The REML analysis of Dataset-1 showed that variances due to genotypes (σ^2^g) were significant for all the traits, indicating the presence of significant variability among genotypes. The range in each trait was high, and high broad heritability (H^2^) > 60% for altogether studied traits were reported in all experiments with a range of 0.72 to 0.91. The mixed model REML analysis of variance revealed a significant variation among the lines for all traits. The Dataset-2 also showed high range of heritability (0.71 to 0.88) for all the five traits (**Table 1**). Best linear unbiased predictors (BLUPs) were obtained for each accession’s traits for both the datasets. The phenotypic and genotypic data for the Dataset-3 was directly taken from the 1K-RiCA^26^.

### Epistasis interaction and DeepMap architecture

The elementary neural network architecture comprises of ‘u’- input variables (I = {g_1_, g_2_, g_3,…,_ g_u_}), ‘h’- hidden layers, and ‘l’- output layers (where, ‘u’, ‘h’ and ‘l’ ∈ N (Set of Natural Numbers)). The genotypic input along with epistasis interactions that include, Additive (A), Dominance (D), Additive × Additive (A × A), Dominance × Dominance (D × D), Additive × Dominance (A × D), are given to the model in the form of tensors/array. The genotypic line is mapped with quantitative/qualitative phenotypic output as,

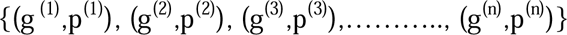

where g^(i)^ is the i^th^-germplasm input containing *g*1 *i*, *g*2 *i*, *g*3 *i*, …, *gn i*} Where ‘n’ is the number of genotypic lines mapped with an associated p^(i)^ phenotypic trait value for ‘m’ number of SNPs germplasm/genotypes. The prediction on the training set is,

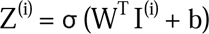

where *Z^(i)^* is the prediction output obtained from an activation function applied on *W^T^* transpose of weighted matrix 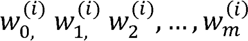, I is a matrix of input containing genotypes and phenotypes for ‘n’ germplasm and ‘m’ SNPs, and b is taken as a bias. The Rectified Linear Unit (ReLU) (Garreta, 2013) activation function is used in hidden layers of DeepMap, and the Linear activation function (Garreta, 2013) is used in the output layer for complex quantitative phenotypes, while sigmoid/SoftMax can be used for qualitative output. The error function for predicted and actual values is subjected to minimize,

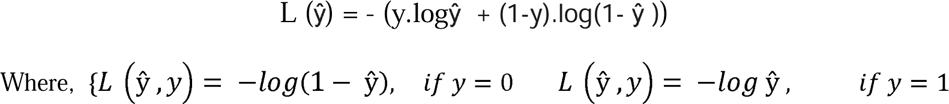

That states if y = 1 then we have to minimize the logŷ to keep ŷ close to y. conversely, if y = 0, then we have to increase log(1- ŷ) given the fact to plummet ŷ. Where *L* is an error function implemented on *y* (actual phenotypic trait value) and ŷ (predicted phenotypic trait value). The combined error function (CEF) for all training examples is,

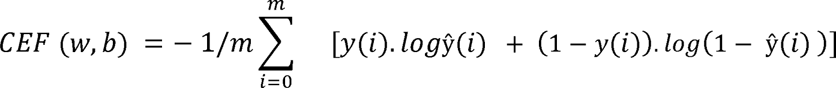

The CEF tends to minimize genotypic patterns through training to the model. The proposed model is based on Dense Neural Networks (DNNs) as shown in **Figure 2**.

### Genotypic input preparation

The genotypic data were pre-processed using R-script provided in the DeepMap package (https://github.com/IRRISouthAsiaHub/DeepMap/) where snpStats (Clayton, 2021), rrBLUP, and sommer (Covarrubias-Pazaran, 2016) were utilized for additive and dominance matrix generation from the plink genotypic data (consists of .bed, .bim, and .fam file formats). This genotypic information is changed into a numeric format: SNPs encoded as ‘0’ represent homozygous allele 2, ‘1’ represents heterozygous allele, and ‘-1’ represents homozygous allele 1. The ‘-1’ was also represented for missing alleles, as the frequency was very low in the dataset. The epistasis interactions are given to the model to train the neural weights and reduce the error loss. After grid search optimization, the DeepMap model has primed with seven hidden layers with 512, 512, 512, 512, 256, 32, and one neural unit, respectively, a single output is anticipated. The functional python programming (Van Rossum, 2010) approach is employed to prevent the shared state, mutable variables and support the construction of pure functions for the model development using open-source libraries, including Keras (Gulli and Pal, 2017), sklearn (Garreta, 2013), scipy (Virtanen et al., 2020), pandas (McKinney, 2010), numpy (Harris et al., 2020) and math.

The core function is main(), which further calls its corresponding subfunctions data_preprocessing(), DeepMap(), plot() and scatter_plot() functions for independent call by value execution. The model is trained on NVIDIA GeForce GTX 1050Ti GPU, having 2200M transistors on 14nm technology with 4096MB memory size configured with CUDA 10.2. The detailed software and hardware information is mentioned in the supplementary. The genotypic data for Datasets-1 and -2 are taken from https://snp-seek.irri.org/ for the 3K rice panel dataset (3380 SNPs). The number of genotypes for both datasets is taken common encompasses 532 for DTF, 1316 for NPT, 1223 for PL, 963 for PH, and 1163 for PY. Each dataset is divided into training and validation set with 4:1 and to uphold the reliability of performance, both sets were split into k-folds cross-validation, where *k* was taken ten (reference). The training set consisted of nine folds, and validation had one-fold.

### Training and evaluation

The architecture of the DNNs used in our experiment is shown in Figure 1. We used ReLU and linear activation functions in the network with the Adam optimizer. The minimum learning rate was set to 0.001 with a factor of 0.5 and patience as 1. The early stopping was placed on validation loss with patience 20. The training set was executed for 10,000 epochs with 2143 batch size on 24GB random access memory. These values are used as a default for the corresponding parameters in our model.

Following the training phase, the model undergoes validation on certain performance parameters, including mean squared error (MSE), root mean squared error (RMSE), R^2^-Error, and scatter plot of normalized predicted vs actual values, and training loss function graph to ensure convergence. The genotyping and phenotyping information for Dataset-3 is available at 1K-RiCA_Data for 353 germplasms and 967 SNPs. The data was trained with default hyper-parameters of DeepMap on the Training set (80%) and validated on the Validation set (20%).

## DATA AVAILABILITY

The genotypic data of 3K Rice can be downloaded from https://snp-seek.irri.org/, and phenotypic data is available at the DeepMap’s GitHub repository. Moreover, the region (Hyderabad and Varanasi) and trait (DTF, NPT, PL, PH, and PY) based models, results, outputs, and validation parameters is also available in the GitHub.

## CODE AVAILABILITY

The DeepMap software has been released to the Python Package Index at https://test.pypi.org/project/DeepMap-1.0/. Its source code and documentation are freely available at https://github.com/IRRISouthAsiaHub/DeepMap, and also in figshare https://figshare.com/articles/software/DeepGS_python_py/23532063. Data pre-processing R-script is available at GitHub repository. Moreover, the region/variety-based models/outputs are available at Result directory of GitHub repository and code for standard GS models is available at Models directory of GitHub repository (including the DeepGS python version).

## ACKNOWLEDGMENTS

The authors express sincere thanks to the Department of Biotechnology (DBT), Government of India for financial support under the project of ‘Development of superior haplotype-based near-isogenic lines (Haplo-NILs) for enhanced genetic gain in rice’ grant (BT/PR32853/AGill/103/1159/2019).

## AUTHOR CONTRIBUTIONS

P.S., and V.K.S. conceived the idea and supervised the study. P.S., V.K.S., A. Kumar and K.T.S. interpreted the results and wrote the first draft of the manuscript. A. Kumar. contributed to the development and evaluation of DeepMap. A.K., K.T.S. and N.G. contributed to the statistical analysis of the genotypic data. U.M.S., C.V. and K.J.P. generated the phenotypic data in IRRI-SAH and ISARC locations. P.J.P. pre-processed and analysed the phenotypic data. P. S., V. K. S., A. Kohli, W.H., B.M. and S.B provided comments on the manuscript and edited the MS. All authors read and approved of the final manuscript.

## ADDITIONAL INFORMATION

The supplementary file for this research manuscript is attached.

## Supporting information

### Supplementary figures

**Figure Legends**

**Figure S1. Training and testing loss for Dataset-1 and Dataset-2.** Training and testing loss for five phenotypic traits of IRRI-SAH (Hyderabad) and ISARC (Varanasi) location iterated for 10,000 epochs (a) training and testing loss converging after 1,433^rd^ epoch on days to flowering (DTF) dataset (b) training and testing loss converging after 148^th^ epoch on NPT dataset (c) training and testing loss converging after 587^th^ epoch on PH dataset (d) training and testing loss converging after 377^th^ epoch on PL dataset e. training and testing loss converging after 3,987^th^ epoch on PY dataset (f) training and testing loss converging after 864^th^ epoch on DTF dataset (g) training and testing loss converging after 206^th^ epoch on NPT dataset (h) training and testing loss converging after 945^th^ epoch on PH dataset (i) training and testing loss converging after 3,488^th^ epoch on PL dataset (j) training and testing loss converging after 6,245^th^ epoch on PY dataset.

**Figure S2. Scatterplot for actual vs predicted values.** The actual and predicted quantitative phenotypic trait value for five traits of two locations (a) for Hyderabad location, DTF trait showed 0.74 correlation on 266 predicted lines, NPT showed 0.65 correlation with 658 lines, PH showed 0.76 correlation with 612 lines, PL showed 0.66 correlation with 266 lines, PY showed 0.70 correlation with 582 lines (b) For Varanasi location, DTF trait showed 0.84 correlation on 266 predicted lines, NPT showed 0.55 correlation with 658 lines, PH showed 0.72 correlation with 612 lines, PL showed 0.60 correlation with 266 lines, PY showed 0.71 correlation with 582 lines.

***Supplementary tables* (**Supplementary tables are in Excel file)

**Table S1.** Input file 1 - Single Nucleotide Polymorphisms (SNPs) for 'n' genotypic lines and 'm' markers

**Table S2.** Input file 2 - Generated additive information for 'n' genotypic lines and 'm' markers.

**Table S3.** Input file 3 - Generated dominance information for 'n' genotypic lines and 'm' markers

**Table S4.** Input file 4 - Phenotypic trait values for 'n' genotypic lines

**Table S5.** Prediction performance of DeepMap for 3K rice panel accessions on days to 50% flowering (DTF) in IRRI-SAH location

**Table S6.** Cross-validation (CV) performance on DeepMap for 3K rice panel accessions on days to 50% flowering (DTF) in IRRI-SAH location

**Table S7.** Prediction performance of DeepMap for 3K rice panel accessions on days to 50% flowering (dtf) in ISARC location

**Table S8.** Cross-validation (CV) performance on DeepMap for 3K rice panel accessions on days to 50% flowering (dtf) in ISARC location

**Table S9.** Prediction performance of DeepMap for 3K rice panel accessions on number of productive tillers (NPT) in IRRI-SAH location

**Table S10.** Cross-validation (CV) performance on DeepMap for 3K rice panel accessions on number of productive tillers (NPT) in IRRI-SAH location

**Table S11.** Prediction performance of DeepMap for 3K rice panel accessions on number of productive tillers (NPT) in ISARC location

**Table S12.** Cross-validation (CV) performance on DeepMap for 3K rice panel accessions on number of productive tillers (NPT) in ISARC location

**Table S13.** Prediction performance of DeepMap for 3K rice panel accessions on plant height (PH) in IRRI-SAH location

**Table S14.** Cross-validation (CV) performance on DeepMap for 3K rice panel accessions on plant height (PH) in IRRI-SAH location

**Table S15.** Prediction performance of DeepMap for 3K rice panel accessions on plant height (PH) in ISARC location

**Table S16.** Cross-validation (CV) performance on DeepMap for 3K rice panel accessions on plant height (PH) in ISARC location

**Table S17.** Prediction performance of DeepMap for 3K rice panel accessions on panicle length (PL) in IRRI-SAH location

**Table S18.** Cross-validation (CV) performance on DeepMap for 3K rice panel accessions on panicle length (PL) in IRRI-SAH location

**Table S19.** Prediction performance of DeepMap for 3K rice panel accessions on panicle length (PL) in ISARC location

**Table S20.** Cross-validation (CV) performance on DeepMap for 3K rice panel accessions on panicle length (PL) in ISARC location

**Table S21.** Prediction performance of DeepMap for 3K rice panel accessions on plot yield (PY) in IRRI-SAH location

**Table S22.** Cross-validation (CV) performance on DeepMap for 3K rice panel accessions on Plot Yield (PY) in IRRI-SAH location

**Table S23.** Prediction performance of DeepMap for 3K rice panel accessions on plot yield (PY) in ISARC location

**Table S24.** Cross-validation (CV) performance on DeepMap for 3K rice panel accessions on plot yield (PY) in ISARC location

**Table S25.** Prediction performance of DeepMap on flowering time (FLW) of 1k-RiCA dataset

**Table S26.** Cross-validation (CV) performance of DeepMap on flowering time (FLW) of 1k- RiCA dataset

**Table S27.** Prediction performance of DeepMap on grain yield (GY) of 1kRiCA dataset

**Table S28.** Cross-validation (CV) performance of DeepMap on grain yield (GY) of 1kRiCA dataset

**Table S29.** Prediction performance of DeepMap on plant height (PH) of 1k-RiCA dataset

**Table S30.** Cross-validation (CV) performance of DeepMap on plant height (PH) of 1k- RiCA dataset

**Table S31.** Validation of DeepMap in other crops (wheat, maize and soyabean)

