## Supplemental document for "DeepMap: A deep learning-based model with a four-line code for prediction-based breeding in crops"

**Full title**

**Running title**

*A deep learning model* for genomic prediction in crops

Ajay Kumar^1,4,#^, Krishna T. Sundaram^1,#^, Niranjani Gnanapragasam^1^, Uma Maheshwar Singh^2^, K. J. Pranesh^1^, Challa Venkateshwarlu^`^, Pronob J. Paul^1^, Waseem Hussain^3^, Sankalp Bhosale^3^, Ajay Kohli^3^, Berta Miro^3^, Vikas Kumar Singh^1,2, *^, Pallavi Sinha^1, *^

^1^International Rice Research Institute (IRRI), South-Asia Hub (SAH), Hyderabad, India

^2^International Rice Research Institute (IRRI), South-Asia Regional Centre (SARC), Varanasi, India

^3^International Rice Research Institute (IRRI), Metro Manila, Philippines

^4^University of Missouri, Columbia, Missouri, USA

^#^Contributed equally

**Supporting information**

***Supplementary figures***

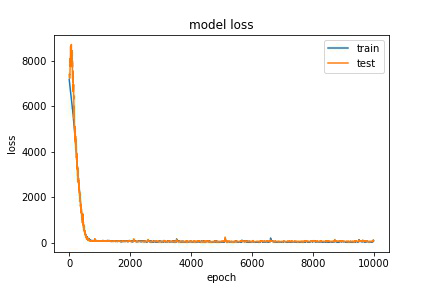

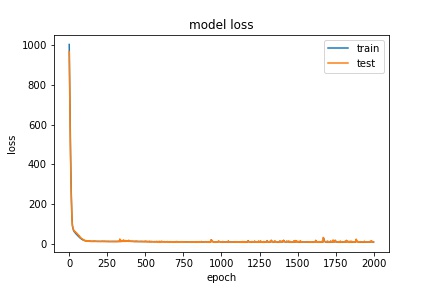

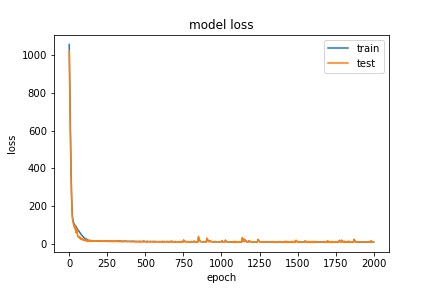

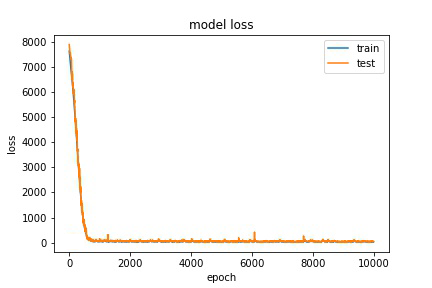
**(a)	(b)**

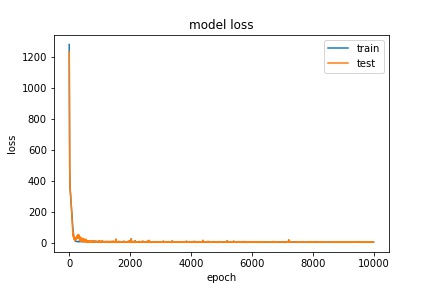

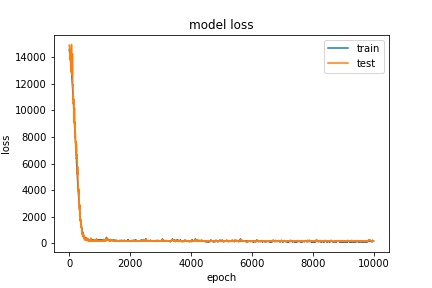

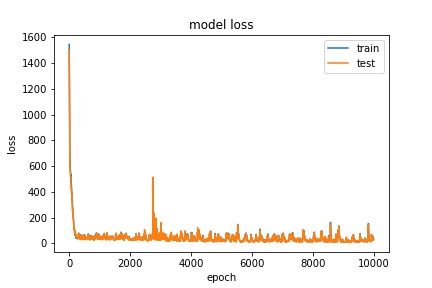

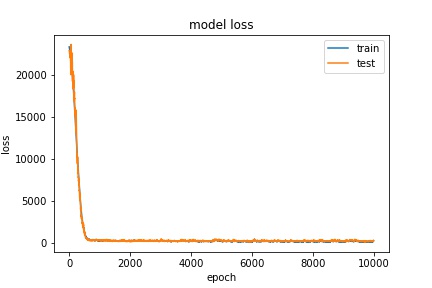

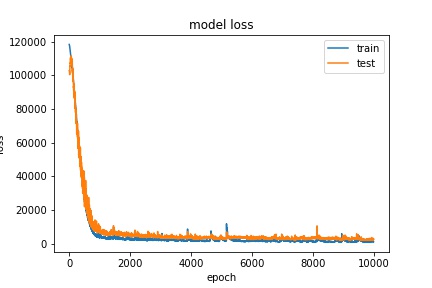

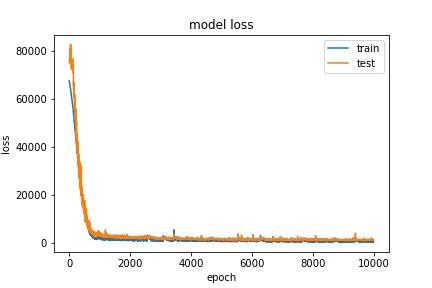

**Dataset-I IRRI-SAH, Hyderabad, India	Dataset-2 ISARC, Varanasi, India**

**Figure S1. Training and testing loss for Dataset-1 and Dataset-2.** Training and testing loss for five phenotypic traits of IRRI-SAH (Hyderabad) and ISARC (Varanasi) location iterated for 10,000 epochs (a) training and testing loss converging after 1,433^rd^ epoch on days to flowering (DTF) dataset (b) training and testing loss converging after 148^th^ epoch on NPT dataset (c) training and testing loss converging after 587^th^ epoch on PH dataset (d) training and testing loss converging after 377^th^ epoch on PL dataset e. training and testing loss converging after 3,987^th^ epoch on PY dataset (f) training and testing loss converging after 864^th^ epoch on DTF dataset (g) training and testing loss converging after 206^th^ epoch on NPT dataset (h) training and testing loss converging after 945^th^ epoch on PH dataset (i) training and testing loss converging after 3,488^th^ epoch on PL dataset (j) training and testing loss converging after 6,245^th^ epoch on PY dataset.

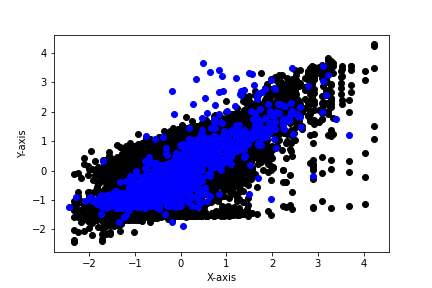

**Actual NPT**

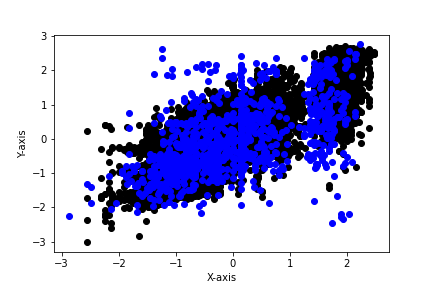

**Actual DTF (50%)**

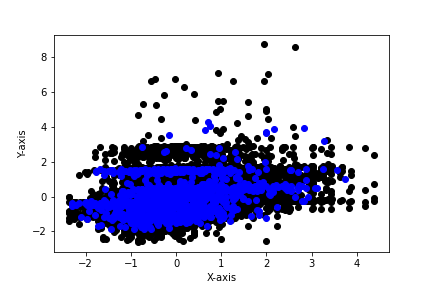

**Actual NPT**

**Predictive NPT**

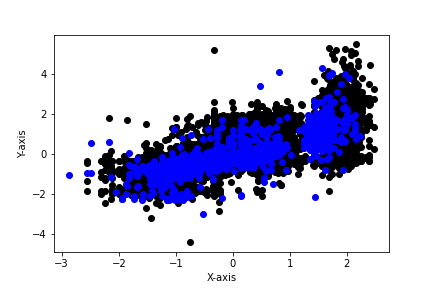

**Actual DTF (50%)**

**Predicted DTF (50%)**

**(b)**

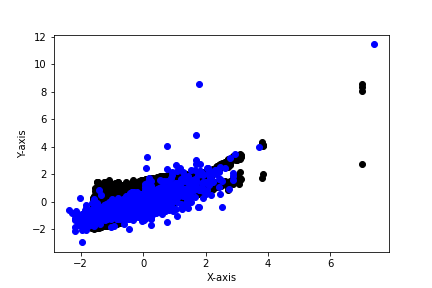

**Actual PY (gram)**

**Predicted PY (gram)**

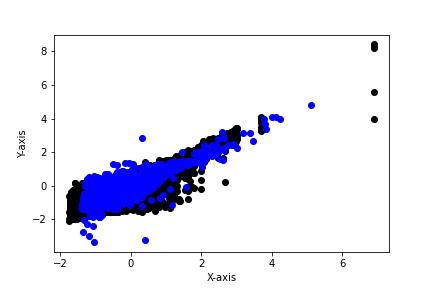

**Predicted PY (gram)**

**Actual PY (gram)**

**Actual PL (cm)**

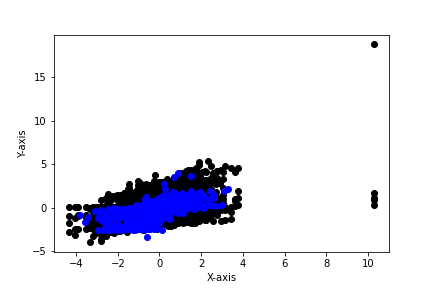

**Predicted PL (cm)**

**Actual PH (cm)**

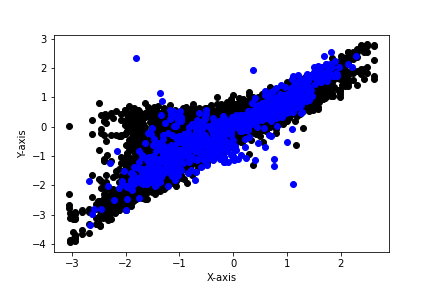

**Predicted PH (cm)**

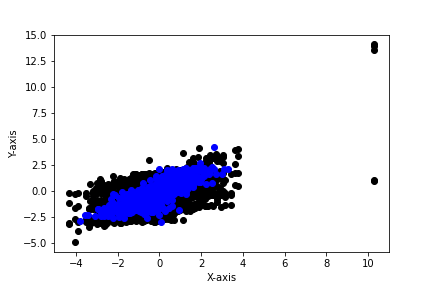

**Predicted PL (cm)**

**Actual PL (cm)**

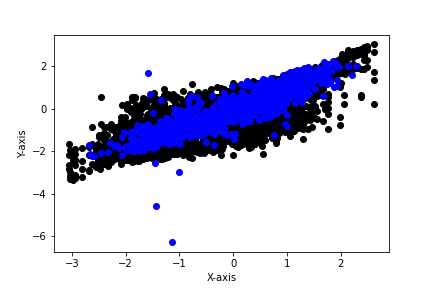

**Actual PH (cm)**

**Predicted PH (cm)**

**Predictive NPT**

**Predicted DTF (50%)**

**Dataset-I IRRI-SAH, Hyderabad, India	Dataset-2 ISARC, Varanasi, India**

**Varanasi, India**

**Predicted -**

**Actual	–**

**(a)**

**Hyderabad, India**

**Figure S2. Scatterplot for actual vs predicted values.** The actual and predicted quantitative phenotypic trait value for five traits of two locations (a) for Hyderabad location, DTF trait showed 0.74 correlation on 266 predicted lines, NPT showed 0.65 correlation with 658 lines, PH showed 0.76 correlation with 612 lines, PL showed 0.66 correlation with 266 lines, PY showed 0.70 correlation with 582 lines (b) For Varanasi location, DTF trait showed 0.84 correlation on 266 predicted lines, NPT showed 0.55 correlation with 658 lines, PH showed 0.72 correlation with 612 lines, PL showed 0.60 correlation with 266 lines, PY showed 0.71 correlation with 582 lines.
